# Optimizing nucleic acid extraction and transcriptome evaluation from low-input, fixed clinical samples

**DOI:** 10.1101/2020.06.02.122150

**Authors:** Katie M. Campbell, Egmidio Medina, Ignacio Baselga Carretero, Yaroslav Teper, Rangasamy Elumalai, Raymond Miller, Kasturi Pal, Jessica Pitzer, Eugenio Daviso, Carsten Carstens, James Laugharn, Willy Hugo, Jia Ming Chen, Agustin Vega-Crespo, Sameeha Jilani, Ivan Perez Garcilazo, Ling Dong, Xinmin Li, Siwen Hu-Lieskovan, Antoni Ribas

**Affiliations:** Department of Medicine, Division of Hematology-Oncology, University of California, Los Angeles (UCLA), Los Angeles, CA; Agilent Technologies, Santa Clara, CA; Covaris, Inc., Woburn, MA; Parker Institute for Cancer Immunotherapy Center at UCLA, Los Angeles, CA; Department of Pathology and Laboratory Medicine, UCLA, Los Angeles, CA; Jonsson Comprehensive Cancer Center; Salt Lake City, UT; Huntsman Cancer Institute, University of Utah, Salt Lake City, UT; Department of Surgery, Division of Surgical Oncology, UCLA, Los Angeles, CA; Department of Molecular and Medical Pharmacology, UCLA, Los Angeles, CA

## Abstract

Tumor biopsies are commonly formalin-fixed and paraffin-embedded (FFPE) for long-term and efficient storage. However, FFPE preservation can greatly compromise the quality of samples, the extraction of nucleic acids, and feasibility of downstream studies, including RNA sequencing. These challenges are especially evident in the studies of clinical trial samples, where the sizes of biopsies often limit the amount of material available for study. Here, we evaluate two nucleic acid extraction kits (Covaris truXTRAC FFPE tNA Plus, QIAGEN miRNeasy FFPE) and three hybridized-capture-based RNA sequencing library preparations (Agilent SureSelect XT RNA Direct, Agilent SureSelect XT HS, and Illumina TruSeq RNA Exome) to evaluate the impact of these sample processing steps on transcriptome evaluation in a melanoma biopsy procured in a clinical setting and preserved by FFPE. While there exist many options for extraction and library preparation that are appropriate for RNA sequencing from FFPE samples, we observed that combinations of experimental approaches may have subtle impacts on downstream analysis, including gene expression quantification and fusion detection.

## Introduction

Biospecimen banking of formalin-fixed, paraffin-embedded (FFPE) solid tumor samples is a common option for long-term storage of clinical samples. While this preservation technique is highly economical, formalin-fixation can greatly affect the quantity and quality of nucleic acids through degradation and chemical modification, complicating the extraction of high-quality nucleic acids from FFPE material(1). Some of these challenges can be addressed by the use of proteinase K to completely solubilize tissues, limiting the fixation time, or introducing heating prior to hybridization(1,2). Regardless, the quality and quantity of extracted material is highly dependent upon the amount of starting material and extraction methodology(3,4).

Reduced quality and quantity of nucleic acids also complicates downstream next generation sequencing analysis(5), particularly in transcriptome profiling, since inefficient or inconsistent sequencing of transcripts can impact the gene expression profile of the sample. FFPE can cause degradation of RNA molecules, resulting in loss of the poly-adenylated (poly-A) tails, which is the target of oligo-dT primers used in poly-A enrichment RNA sequencing strategies. Furthermore, adenine undergoes the most frequent chemical modification by formalin fixation, which further impedes the amplification of poly-A transcripts(2). Thus, ribosomal depletion and hybridized capture approaches have become more prevalent for degraded types of RNA. Both ribosomal depletion and hybridized capture work by enriching the library for transcripts not associated with ribosomal RNA, which dominate the pool of transcripts extracted from a sample. Ribosomal depletion protocols have been described as technically challenging and inconsistent, may exhibit variable performance based upon the level of degradation(6), require a minimum of 400ng of total RNA input, according to standardized methods(1), and result in libraries dominated by highly expressed transcripts. Hybridized capture approaches implement gene-targeting capture probe sets to select transcripts associated with known genes and are able to reduce the difference between lowly and highly expressed transcripts.

Studies in melanoma and other tumor types are further complicated by the size of the lesion or biopsy obtained for study, and the size of the sample can greatly limit the types and number of correlative studies that can be performed. In order to address the criticality of extraction, RNA sequencing library preparations, and the combination of these approaches, we compared two nucleic acid extraction approaches in combination with three hybridized capture-based, RNAseq library preparation protocols to optimize a workflow for transcriptome profiling of clinical FFPE samples.

## Materials and Methods

### Sample overview

The tumor sample (PT0017bx) used in this study was obtained with patient consent through UCLA IRB#16-000029 and preserved by FFPE. Scrolls (4um thick) were procured from the FFPE block by the Tissue and Pathology Core Laboratory at UCLA. The FFPE Kidney and Lung samples used for the nucleic acid extraction optimization were provided by Covaris.

### Nucleic acid extraction

RNA extraction using the QIAGEN miRNeasy FFPE kit was performed according to the manufacturer’s protocol, with one exception. After the first elution was performed, by passing 14 ul of RNase-free water through the RNA elution column, elution was performed two more times, for a total of three elutions. Each elution was preserved individually, to maximize the amount of RNA isolated and assess the added benefit of collecting multiple elutions.

DNA and RNA extraction using the Covaris truXTRAC FFPE tNA Plus (Column) kit, according to the manufacturer’s protocol, specifically on the Covaris E220 evolution platform. Of note, no changes were made to the protocol; specifically, the isopropanol concentration (31.1%) was not altered to impact RNA yield or quality.

RNA quality and quantity were assessed using the Agilent Bioanalyzer 2100 and the RNA Nano chip. The DV200 value was quantified using the Agilent 2100 Bioanalyzer Expert Software, calculating the percent of material that was at least 200nt in length. RNA quality was also assessed by Nanodrop.

Genomic DNA quality and quantity were assessed using the Agilent TapeStation 2200 with the Genomic DNA tape. The DNA integrity number (DIN) was calculated using the 2200 TapeStation Software.

### Library preparation

The Agilent Universal Human Reference (UHR) RNA was used as a positive control for all library preparation approaches.

Agilent SureSelect XT HS workflow was performed with input quantities of 20ng, 50ng, or 100ng of RNA, and libraries were prepared according to a preliminary protocol provided by Agilent. Agilent SureSelect XT RNA Direct was performed with input quantities of 100ng or 200ng of RNA, and libraries were prepared according to the manufacturer’s protocol. For both Agilent kits, hybridized capture was performed using the Agilent SureSelect Human All Exon V7 Exome.

Illumina TruSeq RNA Exome was performed with input quantities of 20ng, 50ng, or 100ng of RNA, and libraries were prepared according to the manufacturer’s protocol. During library preparation, four libraries were pooled prior to the hybridized capture step (PT0017_Qiagen_100ng_Illumina, PT0017_Covaris_50ng_Illumina, PT0017_Covaris_100ng-Illumina, and Agilent_UHR_50ng_Illumina).

Libraries were assessed during preparation using the Agilent Bioanalyzer 2100. Pre-capture libraries were evaluated using the DNA 1000 chip and quantified by selecting for molecules that were 175-700 nucleotides long. Pre-capture libraries with less than 100 ng of double-stranded cDNA (n=5; PT0017_Qiagen_20ng_XTHS, PT0017_Covaris_20ng_XTHS, PT0017_Qiagen_20ng_Illumina, PT0017_Covaris_20ng_Illumina, Agilent_UHR_20ng_Illumina; see **Supplementary Table 2**) were not processed through hybridization and sequencing. If there was not 200ng of dsDNA of pre-capture library available, the entire quantity was used for hybridization (see **Supplementary Table 2**). Post-capture libraries were evaluated using the High Sensitivity DNA chip and quantified by selecting for molecules that were 200-1000 nucleotides long.

### RNA sequencing and analysis

Libraries were pooled by their associated library preparation protocol. Each pool (2nM per library; Agilent SureSelect XT HS, n=6; Agilent SureSelect XT RNA Direct, n=6; Illumina TruSeq RNA Exome, n=5) was sequenced on one lane on the Illumina HiSeq 3000 platform in the Technology Center for Genomics and Bioinformatics at UCLA. Sequencing reads were aligned to the human reference genome (GRCh38) using Hisat2(7), specifically using the reference genome index for GRCh38 provided at ftp://ftp.ccb.jhu.edu/pub/infphilo/hisat2/data/hisat2_20181025.tar.gz. Gene expression was quantified using HTSeq-count(8) and Ensembl v94(9). Gene expression was normalized by library size using the ‘regularized log’ (rlog) transformation and summarized using DESeq2(10). The rlog gene expression values were scaled by gene (i.e. z-score across samples) for visualization purposes (denoted as ‘relative expression’).

Correlative analysis across sequenced libraries was evaluated by identifying the intersection of genes (annotated by Ensembl v94) covered by both the Illumina TruSeq RNA Exome and the Agilent SureSelect Human All Exon V7 Exome probes. Of note, the bed file provided by Illumina (https://support.illumina.com/content/dam/illumina-support/documents/downloads/productfiles/truseq/truseq-rna-exome-targeted-regions-manifest-v1-2-bed.zip) had to be lifted over using the UCSC LiftOver tool (https://genome.ucsc.edu/cgi-bin/hgLiftOver) from GRCh37/hg19 to GRCh38/hg38. Bed files, and the annotated gene targets, were compared using the GenomicRanges R package(11).

Fusion detection was performed by kallisto(12) followed by pizzly(13). Putative fusions were filtered to those with paircount>0 and splitcount>3. Fusions were filtered by genomic position (to remove the presence of multiple annotations).

## Results

### Experimental Overview

We designed a series of experiments to compare the effect of RNA extraction, library preparation, and the combination of these approaches from FFPE tissue on downstream gene expression analysis (**Figure 1**). A tumor sample (PT0017bx) was chosen for having substantial material to perform multiple nucleic acid extractions and multiple RNA sequencing library preparations. Scrolls were obtained in 4um cuts from the FFPE block containing the tumor sample. The first 10 scrolls (40um) were subjected to RNA extraction using the QIAGEN miRNeasy FFPE kit. The next 6 scrolls were grouped into three sets of 2 scrolls (8um of tissue in each set) for DNA and RNA extraction by the Covaris truXTRAC tNA FFPE Plus kit. Libraries were prepared with varied input quantities from both the Covaris-extracted RNA from PT0017bx, QIAGEN-extracted RNA from PT0017bx, and the Agilent Universal Human Reference (UHR) RNA using the Agilent SureSelect XT HS RNA Alpha (20ng, 50ng, 100ng), Agilent SureSelect XT RNA Direct (100ng, 200ng), and the Illumina TruSeq RNA Exome (20ng, 50ng, 100ng).

**Figure 1.**
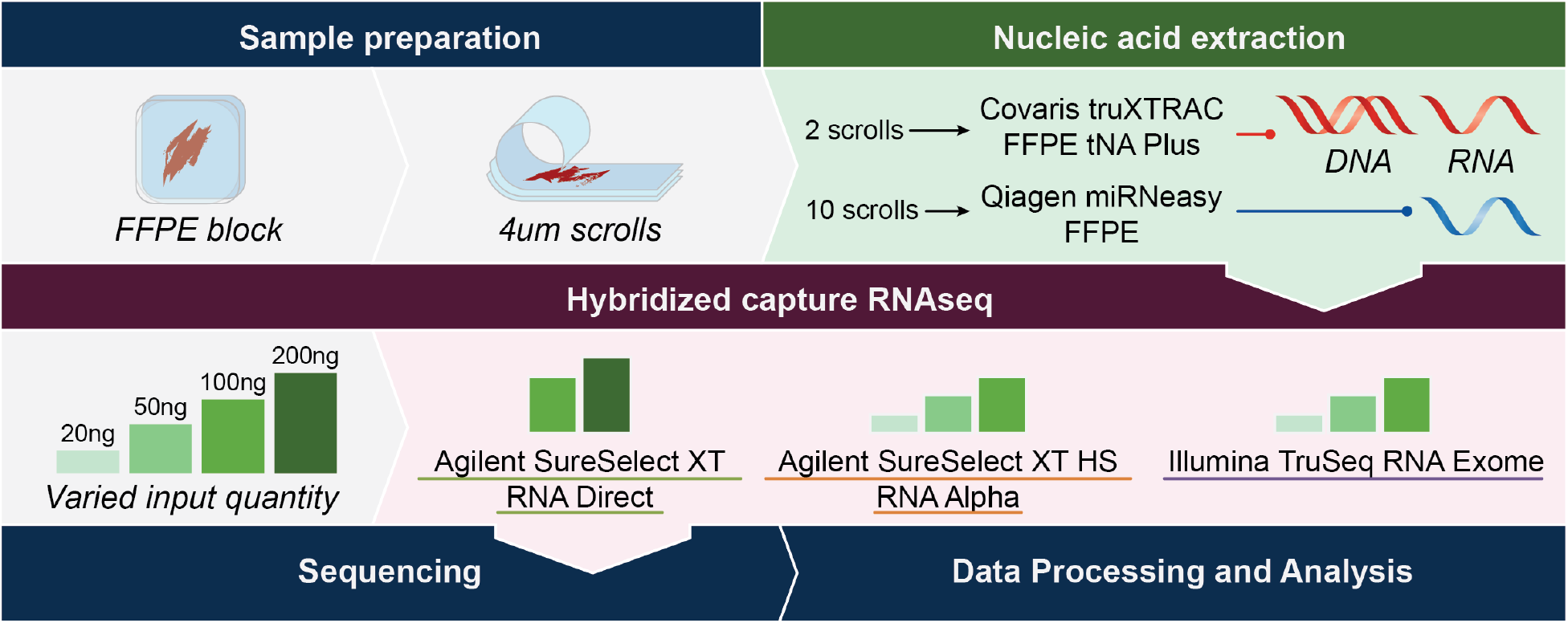
Overview of experiments and analysis performed in this study. Graphic describing the workflow described in this study, including the experimental techniques evaluated and details regarding the variable inputs for each protocol.

### Maximizing the DNA and RNA yield from low-input FFPE tissue

Total RNA was quantified by the Agilent Bioanalyzer (kit), and the purity of the nucleic acids were assessed by Nanodrop (**Supplementary Table S1**, **Figure 2**). There was a total of 33.7ug of RNA extracted from 24um of tissue by the Covaris protocol (6.8-13.5ug per 8um of tissue; 1.4ug RNA/um tissue), and there was a total of 28.9ug of RNA extracted from 40um of tissue by the QIAGEN protocol (0.63-24ug across three elutions; 0.72ug RNA/um tissue; **Figure 2A**). The RNA extracted by Covaris had an average 260:280 ratio of 1.99 (1.98-1.99 across replicates); the RNA extracted by QIAGEN had an average 260:280 ratio of 1.88 (1.76-1.97 across elutions; **Figure 2B**). The average RNA integrity number (RIN) was 2.8 (2.1-3.6 across replicates) in RNA extracted using the Covaris protocol and 1.8 (1-2.3 across elutions) using the QIAGEN protocol. However, RIN values have been shown to not be a sensitive metric for RNA quality in FFPE tissues(14). The DV_200_ (percentage of RNA molecules at least 200 nucleotides [nt] long), which has been shown to be a better indicator of RNA quality, was higher in RNA extracted using the Covaris protocol (average 71.7%; 69-74% across replicates), compared to the QIAGEN protocol (average 58.3%; 57-60% across elutions). Moreover, there was more RNA extracted that was at least 200nt in length using the Covaris protocol (total 24.3ug of RNA; 4.7-9.8ug across replicates; 1.01ug RNA >200 nt per um tissue), compared to the QIAGEN protocol (total 16.5ug of RNA; 0.4-13.7 ng across elutions; 0.41ug RNA >200 nt per um tissue). Overall, the RNA obtained using the Covaris extraction method showed increased yield and longer RNA molecules, compared to the QIAGEN extraction method.

**Figure 2.**
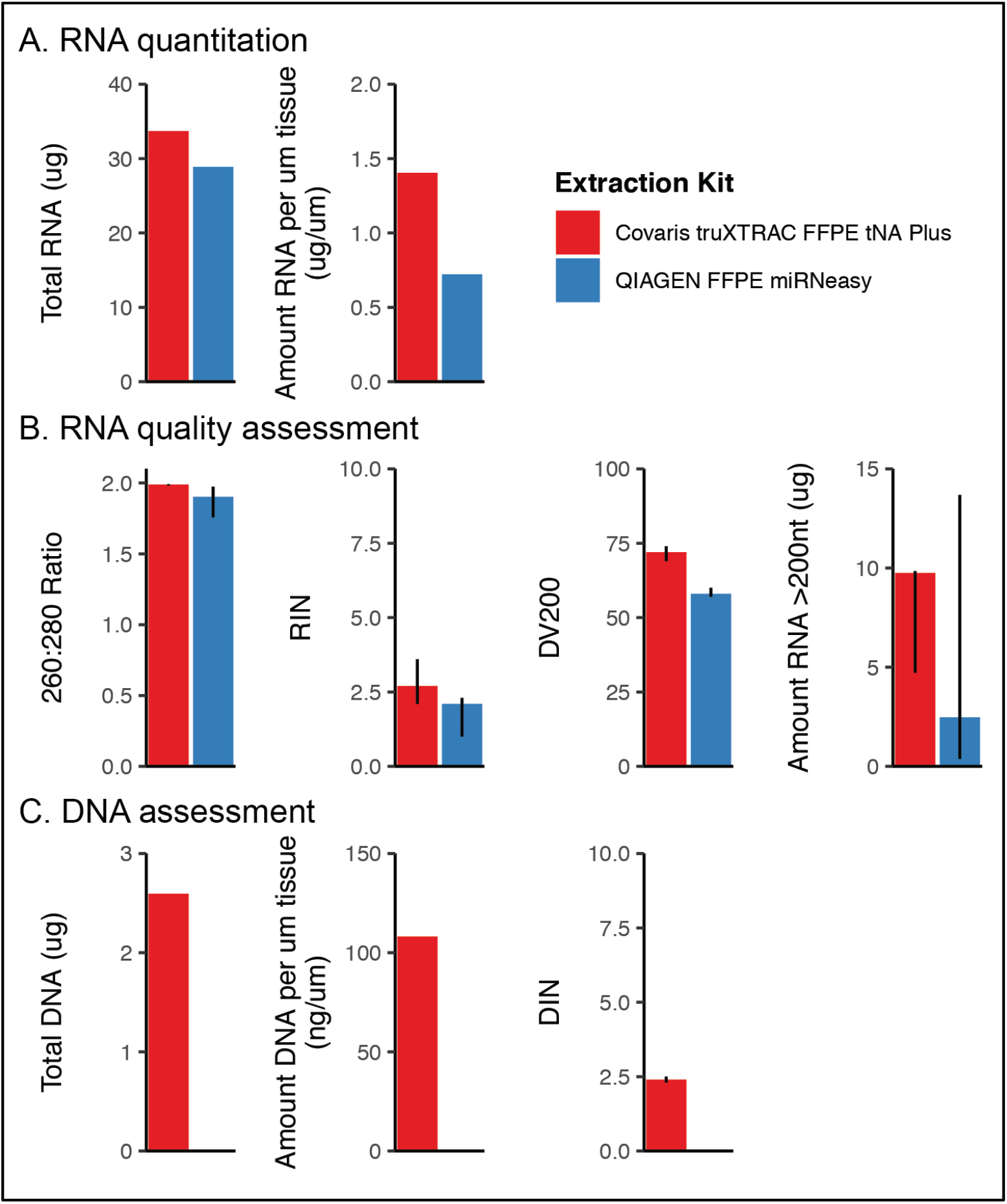
Nucleic acid extraction quantitation and quality assessment. RNA and/or DNA extracted by either the Covaris truXTRAC FFPE tNA Plus kit (red) or the QIAGEN FFPE miRNeasy kit (blue) were evaluated by several measures. (A) RNA was quantified and summarized by the total amount (ug) extracted across triplicates (Covaris) or three elutions (QIAGEN). This value was also summarized by the amount of nucleic acid extracted per micron (um) of tissue. (B) RNA quality was assessed for purity (260:280 ratio) and quality (RNA integrity number [RIN], DV200). Error bars represent the range of values across Covaris replicates (n=3) or the three elutions from the QIAGEN protocol. (C) DNA quantity was summarized by the total amount (ug) extracted across triplicates in the Covaris protocol and the amount per micron of tissue extracted. The DNA integrity number (DIN) was used to summarize quality.

There was a total of 2.60 ng of genomic DNA obtained (0.83-0.93 ng per 8um of tissue; 0.11ng of DNA/um of tissue; **Figure 2B**). The average DNA integrity number (DIN)(15) was 2.4 (2.3-2.5 across replicates). The QIAGEN miRNeasy FFPE kit was only used to obtain RNA, so there was no concurrent DNA extracted from these tissues to evaluate.

We performed extractions in triplicate using the Covaris protocol (8um of tissue per replicate), but only completed a single extraction using the QIAGEN kit (40um of tissue). Thus, to verify that the increased extraction quantities and qualities were not an outlier in the PT0017bx sample, extractions were performed from two additional FFPE samples (kidney and liver tissue, each in triplicate). RNA extracted using the Covaris truXTRAC FFPE tNA Plus kit showed higher yield, increased DV_200_ values, and increased yield of RNA at least 200nt in length, compared to the QIAGEN miRNeasy FFPE Kit in both samples (**Supplementary Table S1**, **Supplementary Figure S1**). Furthermore, the variance in quantity and quality of RNA was reduced using the Covaris kit, compared to QIAGEN for both samples, suggesting increased experimental consistency across these metrics associated with the Covaris-extracted samples.

### Comparing exome capture approaches for transcriptome profiling from FFPE tissues

Previous studies have described the implementation of hybridized capture probes into RNA sequencing library preparation to enrich the sequencing library for RNA fragments transcribed from known gene-encoding regions of the genome(16,17). This is advantageous, since exome reagents contain probes that span the length of the transcript. Conversely, FFPE preservation may cause degradation of the polyadenylated tail of the full-length transcript, preventing transcript selection using traditional poly-A enrichment RNA sequencing methods. We explored three different exome capture-based RNA sequencing approaches to evaluate the consistency and efficacy of these approaches on FFPE-derived RNA: Agilent SureSelect XT HS RNA Alpha, Agilent SureSelect XT HS RNA Direct, and Illumina TruSeq RNA Exome. We examined various input quantities of RNA (**Figure 1**), based upon each protocol’s specifications, that were isolated by either the Covaris or QIAGEN protocol. Agilent Universal Human RNA (UHR) was used as a positive control across these protocols.

All three library preparation protocols had similar workflows (**Figure 3**): fragmentation of RNA, cDNA synthesis, PCR amplification of cDNA, hybridized capture of cDNA using probes to the gene-coding exome, and PCR amplification of post-capture libraries. One step to highlight in the Illumina workflow was the pooling of 4 samples prior to the hybridized capture; other samples were not pooled until directly prior to sequencing.

**Figure 3.**
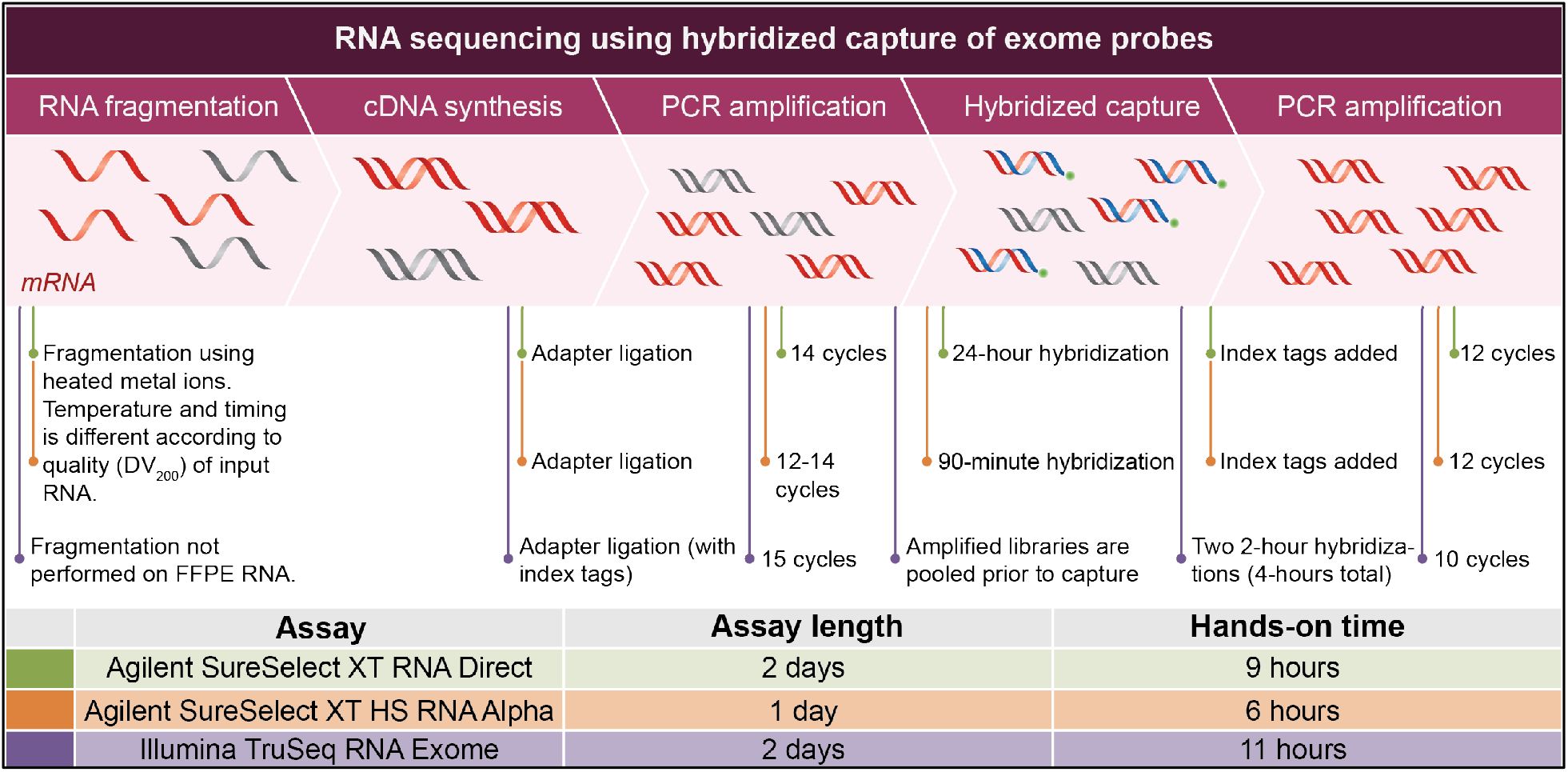
Overview of hybridized capture RNA sequencing library preparation protocols. Graphic describing the sequence of steps associated with the library preparation protocols evaluated in this study. Differences between the protocols are indicated by the comments below (denoted by color).

Libraries were assessed pre- and post-hybridized capture in all three protocols. The amount of dsDNA following the first PCR amplification was loosely correlated with input quantity across library preparations (R^2^=0.46, p=5.6e-4; **Supplementary Figure S2**). Pre-capture libraries with less than 100ng of material were not processed through hybridization and sequencing (see **Methods** and **Supplementary Table S1**).

Post-capture libraries were assessed on the Agilent Bioanalyzer 2100 (High-sensitivity DNA chip). The average size of post-capture libraries was higher in those prepared using either the Agilent SureSelectXTHS RNA Alpha (average 400bp long; 363-444bp) or Illumina TruSeq RNA Exome (average 381bp long; 378-392) protocols, compared to the Agilent SureSelectXT RNA Direct (average 305bp long; 290-319bp) protocol (**Figure 4A-B**, **Supplemental Table S2**). In particular, the combination of the Covaris RNA extraction with the Agilent SureSelectXTHS library preparation resulted in the amplification and capture of longer RNA fragments (average 438bp; 432-444bp). Libraries were pooled, according to their protocols, and each pool (containing 5-6 libraries) was sequenced on one lane of the Illumina HiSeq3000 platform to a depth of 44-207 million (M) reads (average 139M reads; **Figure 4C**). The number of reads per library was most highly variable across libraries prepared using the Illumina TruSeq protocol (44-207M reads; standard deviation of 73.5M reads), particularly across the libraries that were pooled prior to hybridization. However, alignment rates still ranged from 94.54-98.06% (average 96.64%; **Figure 4D**) across all combinations of extractions and library preparations.

**Figure 4.**
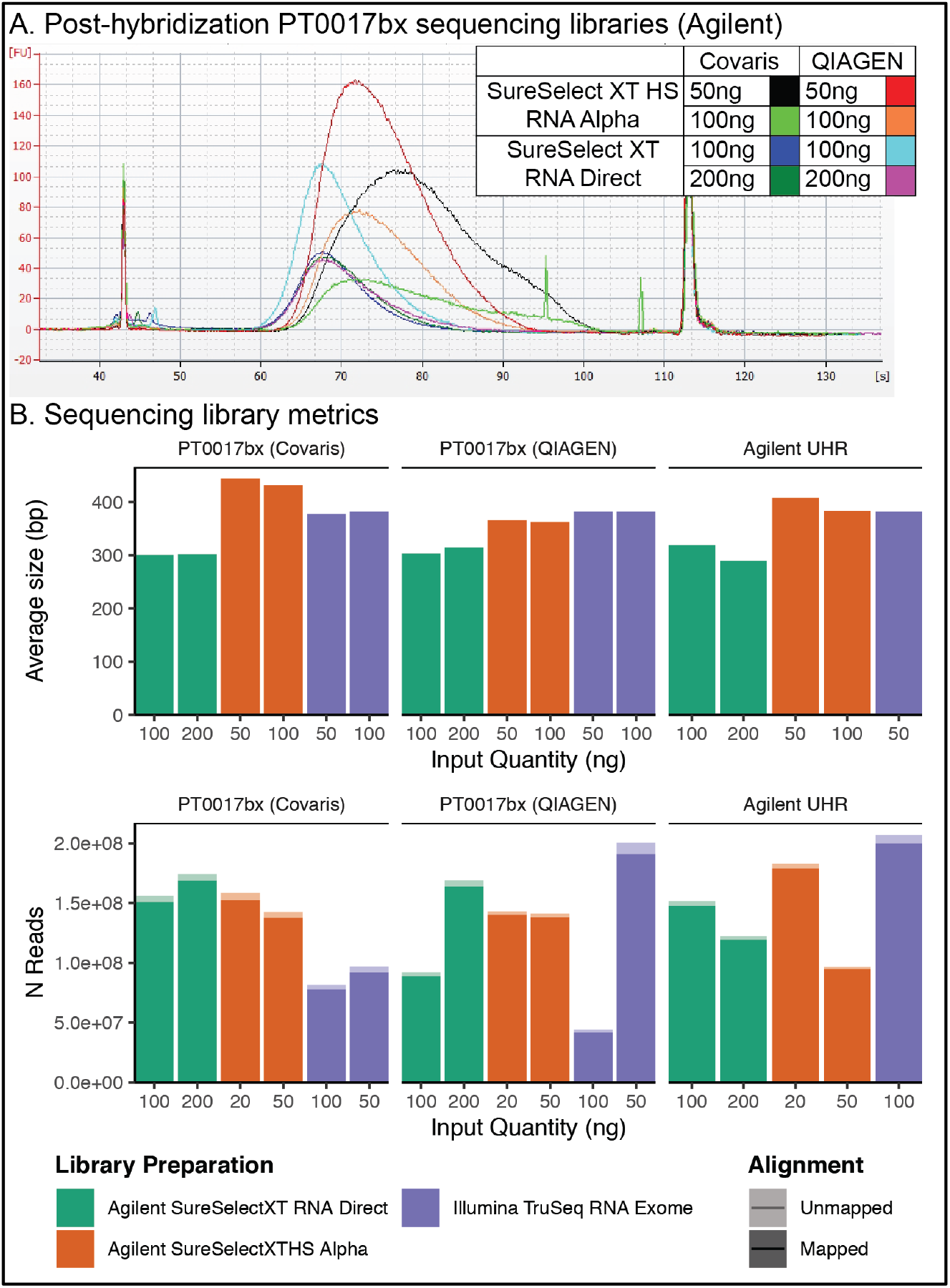
Sequencing and alignment metrics. A. Representative traces from the Agilent Bioanalyzer (High Sensitivity DNA chip) assessment of post-hybridization libraries prepared from the PT0017bx RNA (either Covaris or QIAGEN) and either the Agilent SureSelect XT RNA Direct or XT HS RNA Alpha library preparation kits. B. The average size of dsDNA molecules (top panel; y-axis) was quantified by the Agilent Bioanalyzer for each library (x-axis), and the total number of reads generated for each library (bottom panel; y-axis) was quantified using samtools flagstat tool. The fill color of each bar indicates the library preparation kit; each input quantity is separated along the x-axis; the transparency of the bars for the number of reads generated shows aligned and unaligned reads.

### Evaluating the impact of extraction and library preparation methods on gene expression analysis

We hypothesized that the quality of the input RNA and the library preparation approach may impact subsequent gene expression analysis. Overall, both the Pearson correlation and coefficient of determination between PT0017bx libraries was very high (r=0.949-1.00; median 0.979; r^2^=0.953-1.00; median 0.985; **Figure 5A**). However, principal component analysis (PCA) revealed distinct clustering based upon library preparation manufacturer (PC1) and the combination of extraction method and library preparation (PC2; **Figure 5B**).

**Figure 5.**
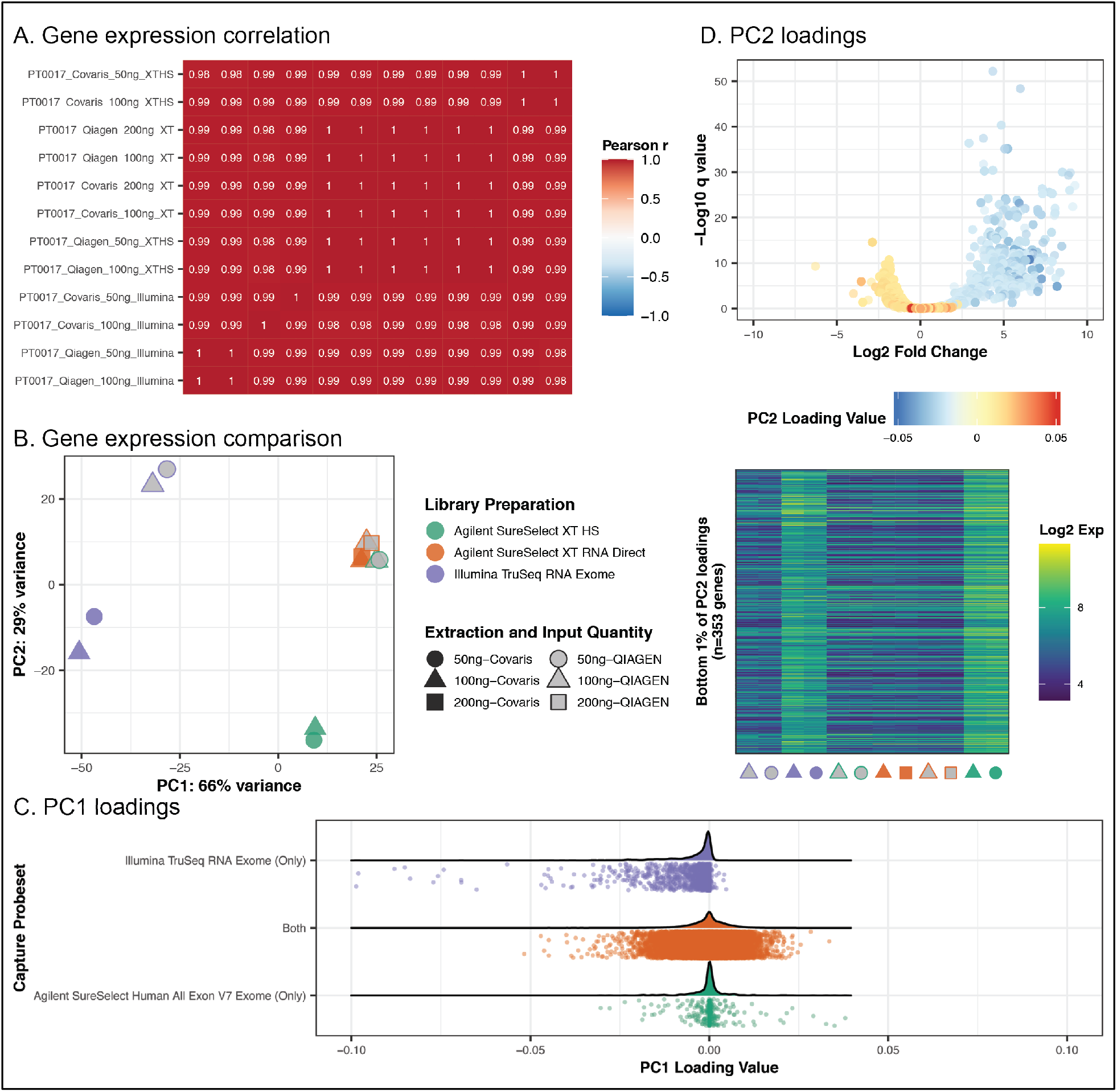
Gene expression comparison. A. Correlation of gene expression (library size-normalized counts across all genes) between libraries. B. PCA plot showing the clustering of points based upon library preparation (color/outline), input quantity (shape), and extraction protocol (grey vs. opaque). C. PC1 loading values of individual genes (x-axis) are shown as both density and point plots. Points are colored based upon whether the gene falls within the targeted genomic region of either Agilent, Illumina, or both exome capture probesets. (D) Differential expression analysis was performed on PC2-low (Covaris-extracted RNA and AgilentSureSelect XT HS or Illumina TruSeq RNA Exome libraries) versus PC2-high (QIAGEN-extracted RNA and/or Agilent SureSelect XT RNA Direct libraries). Genes were filtered to the genes in the top and bottom 1% of PC2 loadings (from panel B; n=706 genes). The volcano plot shows the q value, Log2-fold change (x-axis), comparing PC2-low versus PC2-high, and the PC2 loading value of the corresponding points. The heatmap shows the Log2-library-normalized expression of the genes in the bottom 1% of PC2 loadings (n=353; y-axis) per library (x-axis, indicated by the color/point shown in panel B).

Since samples clustered by the manufacturer along PC1, we evaluated the genomic regions and associated genes targeted by each capture reagent. There were 24,491 genes associated with the regions targeted by the Agilent SureSelect Exon V7 reagent, and 26,951 genes associated with the regions targeted by the Illumina TruSeq RNA Exome probe set. There were 24,101 genes in common between these two probe sets. The factor loadings of the genes uniquely associated with the Illumina TruSeq RNA Exome probe set (n=3,229) were lower on PC1 (Wilcox test p<0.01), revealing that the genes uniquely targeted by the Illumina probe set were contributing to slight differences in the global expression profile (**Figure 5C**).

The defined probe sets were not sufficient to describe differences along PC2. There appeared to be distinct differences by combining the Covaris extraction technique with either the Illumina TruSeq RNA Exome or Agilent SureSelect XT HS library preparation approach (lower PC2 coordinates), but not the Agilent SureSelect RNA Direct protocol. Differential expression analysis was performed, comparing the PT0017, Covaris-extracted RNA with the Agilent SureSelect XT HS or Illumina TruSeq library preparations (PC2-low) to the QIAGEN-extracted RNA and Agilent SureSelect XT RNA Direct library preparations (PC2-high). Genes that had positive PC2 loadings were more highly expressed in PC2-high libraries, compared to PC2-low libraries (**Figure 5D**). The observed average size of molecules in the post-capture libraries and the slightly higher expression of a subset of genes in PC2-low libraries suggested that the combination of Covaris extraction with either the TruSeq or SureSelect XT HS chemistry was able to capture some information lost using the QIAGEN extraction or the SureSelect XT RNA Direct technologies.

### Evaluating fusion detection across library preparation and extraction protocols

Previous studies have described the ability to detect gene-gene fusions from FFPE-derived RNA, particularly the increased ability to detect these fusions using hybridized capture approaches(16). Gene-gene fusions were identified in each library and were compared across each library. The Agilent UHR contains RNA derived from 10 tumor cell lines (18), and previous studies have described fusion events in this reference RNA (19,20). We confirmed a subset of these events in the libraries prepared from these samples, including *GAS6-RASA3, BCAS3-BCAS4, DEPDC1B-ELOVL7*, and *MYO9B-FCHO1.* All of these previously described fusions were detected in the Agilent SureSelect XT HS kits, but only a subset was detected in Agilent SureSelect XT RNA Direct and Illumina RNA Exome kits (**Figure 6**). Of note, the previously observed *ARFGEF2-SULF2* (5/5 libraries), and *SULF2-PRICKLE2* (4/5 libraries) fusions were also present, but were filtered out due to low read support; and the *TMPRSS2-ERG* and *BCR-ABL1* fusions were not detected in any libraries. There were eleven fusions that were only detected in a single library, ten of which were only observed in the Illumina TruSeq RNA Exome library (**Figure 6A**).

**Figure 6.**
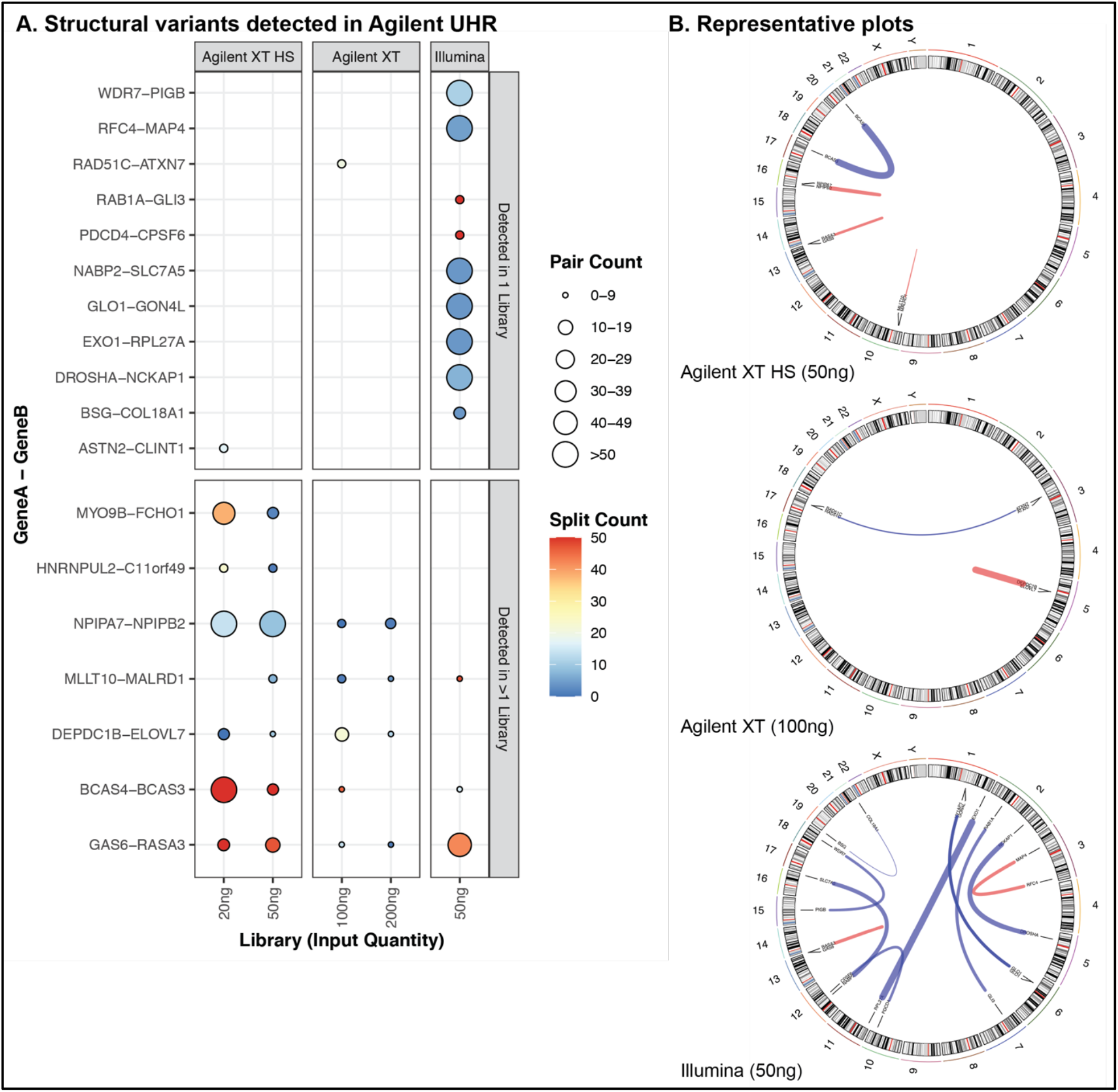
Gene-gene fusions detected in Agilent UHR. Structural variants were detected in each Agilent UHR library, and summarized by the annotated gene-gene fusions. A. The union of fusions detected across all libraries were divided into those that were detected in only one (top panel) or more than one (bottom panel) library. Each point indicates that the fusion was detected in the corresponding library (x-axis) with paired-end reads spanning the gene-gene fusion junction (pair count; size) or reads overlapping and containing the junction (split count; color). Libraries are ordered along the x-axis by input quantity (ng) and grouped by library preparation kit. B. Fusions in A were sorted to those with a pair count >0 and a split count >3 within each sample, and a subset (one library per preparation kit) displayed as circos plots. Fusions in blue show interchromosomal rearrangements, and fusions in red indicate intrachromosomal rearrangement.

We used the same fusion detection strategy on libraries derived from PT0017bx (**Supplementary Figure 3**). Fusions have not been described nor validated in this sample previously, so the results were reduced to those detected in more than one library. Of note, the *NAIP-OCLN* fusion, detected in all library preparations and extractions, has been previously described as a chimeric transcript that may be due to internally deleted or truncated versions of these genes in the germline (21,22). None of the other observed fusions are reported as cancer-associated fusions.

## Discussion

The objective of this study was to directly compare several protocols for nucleic acid extraction and subsequent RNA sequencing, specifically from FFPE-preserved tissues, to ultimately identify optimal experimental approaches to study clinical samples. These studies were performed on an FFPE-preserved patient tumor biopsy (PT0017bx), to simulate a clinically relevant scenario, with technical replicates.

Both the Covaris truXTRAC FFPE tNA Plus and QIAGEN miRNeasy FFPE protocols successfully extracted RNA from FFPE-preserved tissue. However, the Covaris protocol demonstrated increased consistency in control samples, higher yields of RNA, and increased average length of RNA molecules. The major difference between these two protocols is the use of the Covaris Adaptive Focused Acoustics technology, using sonication to facilitate emulsification of paraffin and rehydrate the tissue for DNA extraction without using harsh organic solvents, suggesting the negative impact of organic solvents and method of paraffin removal on overall yield of material. We would also like to highlight that QIAGEN offers the AllPrep DNA/RNA FFPE Kit for concurrent extraction of DNA and RNA, and while the AllPrep protocol was not used in this study, technical documentation reports similar yields of material as the miRNeasy FFPE Kit.

RNA integrity is inherently compromised by FFPE preservation, which is reflected by the RIN value. RNA extracted by both Covaris and QIAGEN protocols displayed low RIN values. However, the length of the extracted RNA can greatly affect the downstream analysis, particularly by next generation sequencing. For example, degraded, overly cross-linked, or shorter RNA molecules can attenuate expression profiles or limit detection of full-length transcripts, which limits identification of alternative splicing events or gene-gene fusions. We did observe downstream consequences on library preparation and transcriptome profiling that were associated with the quality of the RNA extracted by either protocol. While the slight differences observed in gene expression may be associated with spatial heterogeneity, the sampling strategy involved obtaining sequential 5um cuts from the same block for the two extraction strategies. However, the gene expression differences across libraries, where slightly higher expression was observed (PC2), were observed in Covaris-extracted samples prepared using either the Agilent SureSelect XT HS and Illumina TruSeq RNA Exome kits, and not in the Agilent SureSelect XT RNA Direct Kit.

The greatest effect of input quantity was on the library quantities following the first round of amplification during library preparation. Only one library with an input quantity of 20ng displayed enough material to proceed to hybridized capture; however, this was only in the case of the Agilent UHR, included as a positive control, during the Agilent SureSelect XT HS library preparation. Based upon the results of this study, we would suggest using at least 50ng of RNA input for library preparation; however, 100ng may be more highly recommended to increase the pool of transcripts sampled and the chances of observing lowly-abundant genes or capturing gene-gene fusions.

With the goal of transcriptional profiling in clinical FFPE samples in mind, we would like to highlight that the differences across the gene expression landscape were very subtle across combinations of extraction and library preparation strategies. We did observe increased extraction and amplification of longer RNA fragments, particularly in libraries prepared with either the Illumina TruSeq RNA Exome or the Agilent SureSelect XT HS protocols on RNA extracted using the Covaris truXTRAC FFPE tNA plus kit. Conclusively, it is important to note that the choice of extraction protocol, library preparation kit, and exome capture reagent may be critical for the study of clinical samples, especially if gene expression profiling is to be used for biomarker evaluation.

**Table 1.** Comparison of nucleic acid extraction techniques. This table describes the qualitative and quantitative differences between the two nucleic acid extraction techniques evaluated in this study.

| Task | QIAGEN miRNeasy FFPE | Covaris truXTRAC tNA FFPE Plus |
| --- | --- | --- |
| Deparaffinization | Solution-based, using the QIAGEN Deparaffinization Solution (Cat No. 19093), which contains organic solvents. | Mechanical, using the Covaris Adaptive Focused Acoustics® (AFA) technology, which focuses ultrasonic waves at the sample in solution, emulsifying the paraffin. |
| Tissue digestion | Proteinase K is used to lyse the sample and release nucleic acids. | Proteinase K is used to lyse the sample and release nucleic acids. |
| Nucleic acid isolation | Total RNA is released following deparaffinization and tissue digestion. Genomic DNA is removed using a DNase step. | Total RNA is released during deparaffinization using lower concentrations of proteinase K, while genomic DNA remains in the pellet (following centrifugation) with the tissue. Genomic DNA is released by rehydrating the tissue, by AFA, and treating the tissue with higher concentrations of Proteinase K. |
| User-based modifications | For low input samples, multiple elutions from the column are collected to maximize RNA yield. | Isopropanol concentrations can be modified to increase either RNA yield or quality (DV200). |
| Procedure time | Turn-around time of 85 minutes (~20 minutes hands-on time) per sample | Turn-around time of 120 minutes (~15 minutes hands-on time; <5 minutes of machine time) per sample |
| Result | Total RNA | Genomic DNA and total RNA |
| Other notes | QIAGEN also offers the QIAGEN AllPrep DNA/RNA FFPE procedure, which can isolate DNA and RNA from the same input sample. This was not explored in this study. Product details describe comparable yields of RNA in the AllPrep kit as the RNA-specific kit. |  |

## Acknowledgements

We would like to thank the Vondriska laboratory at UCLA for the use of their Covaris E220 Evolution for extraction. KMC is supported by the UCLA Tumor Immunology Training Grant (NIH T32CA009120) and the Cancer Research Institute Postdoctoral Fellowship Program. AR is supported by the National Institute of Health (R35 CA197633), the Ressler Family Fund, the Agilent Thought Leader Award, a Stand Up to Cancer-Bristol-Meyer Squibb Catalyst Research Grant (Grant Number: SU2C-AACR-CT06-17). This research grant is administered by the American Association for Cancer Research, the scientific partner of SU2C. AR is a member researcher at the Parker Institute for Cancer Immunotherapy.

## Author Contributions

KMC and AR conceived experiments and supervised the study. KMC, EM, RE, RM, KP, JP, ED, JP, JL, WH, JMC, LD, XL, SHL, and AR designed the experiments. IBC, AVC, SJ, and IPG performed sample processing and oversaw biobanking. KMC, EM, YT, and RE conducted experiments. LD and XL performed sequencing. KMC, EM, CC, and WH analyzed the data.

KMC and EM wrote and revised the manuscript.

