## Supplementary Appendix for "Optimizing nucleic acid extraction and transcriptome evaluation from low-input, fixed clinical samples"

### Supplementary Figures

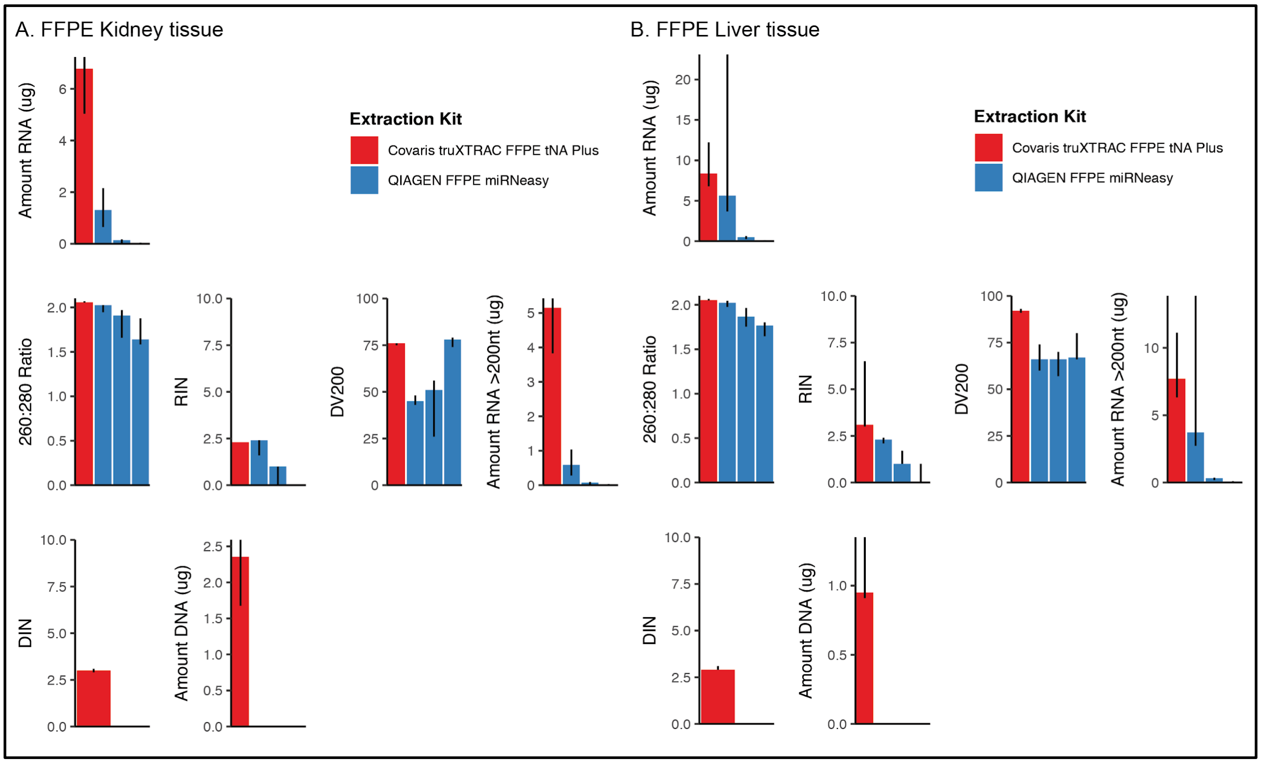


*Supplementary Figure S1. Nucleic acid extraction from additional samples.*

Scrolls were obtained from two additional FFPE samples (see **Methods**), a kidney and lung tissue sample. DNA and/or RNA were extracted from these samples, using either the Covaris truXTRAC FFPE tNA Plus kit (red) or the QIAGEN miRNeasy FFPE kit (blue). (A) RNA quantity was measured by total RNA extracted (ug) and total RNA (ug) extracted per um of tissue. (B) RNA quality was assessed by RIN, 260:280, $DV_{200}$, and amount of RNA >200nt (ug). (C) DNA extracted by the Covaris kit was evaluated by DIN, total DNA extracted (ug), and total DNA (ug) per um of tissue.

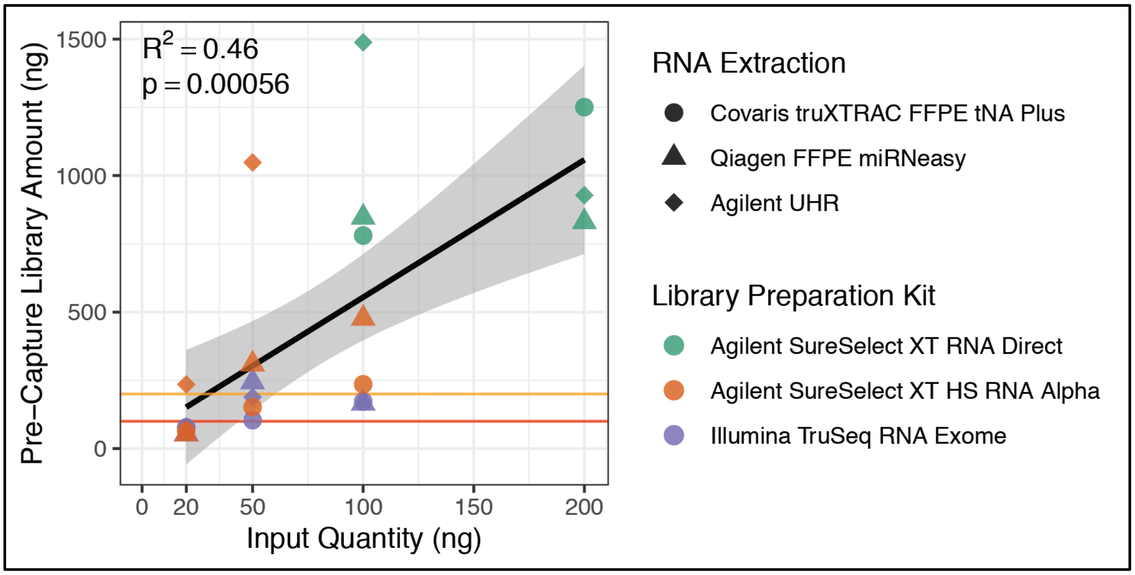


*Supplementary Figure S2. Correlation of pre-capture libraries and input quantity*

Correlation between input quantity for library preparation (ng; x-axis) with the amount of material generated following the first PCR amplification in the library preparation protocol (ng; y-axis). Point shape indicates the starting sample (PT0017 RNA extracted by either Covaris or QIAGEN; Agilent UHR intact RNA). Point colour indicates the corresponding library preparation protocol.

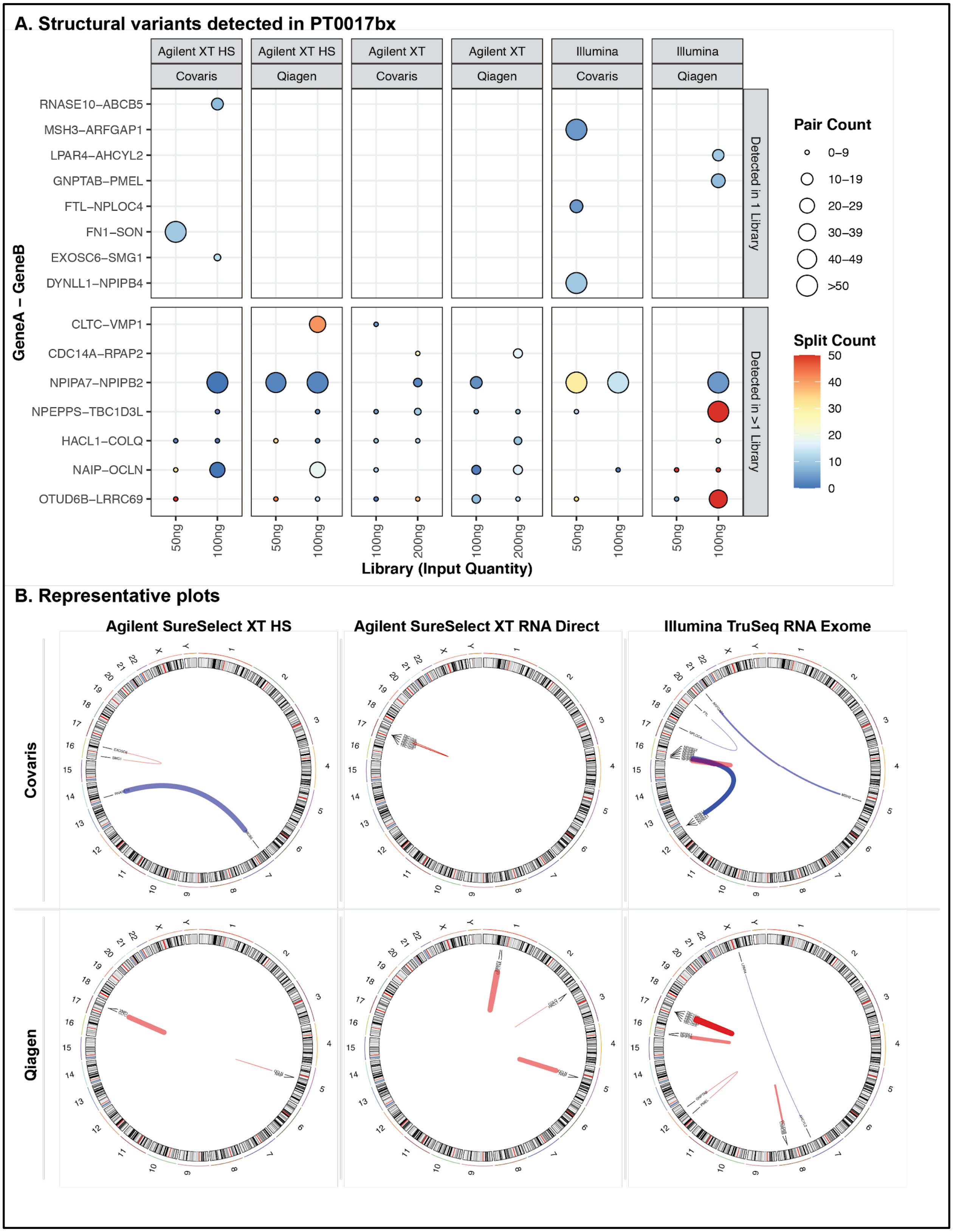
A. Structural variants detected in PT0017bx
Agilent XT HS
Covaris
Agilent XT HS
Qiagen
Agilent XT
Covaris
Agilent XT
Qiagen
Illumina
Covaris
Illumina
Qiagen
GeneA – GeneB
RNASE10–ABCB5
MSH3–ARFGAP1
LPAR4–AHCYL2
GNPTAB–PMEL
FTL–NPLOC4
FN1–SON
EXOSC6–SMG1
DYNLL1–NPIPB4
CLTC–VMP1
CDC14A–RPAP2
NPIPA7–NPIPB2
NPEPPS–TBC1D3L
HACL1–COLQ
NAIP–OCLN
OTUD6B–LRRC69
Detected in 1 Library
Detected in >1 Library
50ng
100ng
50ng
100ng
100ng
200ng
100ng
200ng
50ng
100ng
50ng
100ng
Library (Input Quantity)
Pair Count
0–9
10–19
20–29
30–39
40–49
>50
Split Count
50
40
30
20
10
0
B. Representative plots
Agilent SureSelect XT HS
Agilent SureSelect XT RNA Direct
Illumina TruSeq RNA Exome
Covaris
Qiagen

*Supplementary Figure 3. Structural variants detected in PT0017bx*

Structural variants were detected in each PT0017bx library and summarized by the annotated gene-gene fusions. A. The union of fusions detected across all libraries were divided into those that were detected in only one (top panel) or more than one (bottom panel) library. Each point indicates that the fusion was detected in the corresponding library (x-axis) with paired-end reads spanning the gene-gene fusion junction (pair count; size) or reads overlapping and containing the junction (split count; color). Libraries are ordered along the x-axis by input quantity (ng) and grouped by library preparation kit. B. Fusions in A were sorted to those with a pair count >0 and a split count >3 within each sample, and a subset (one library per extraction [row] and preparation kit [column] combination) displayed as circos plots. Fusions in blue show interchromosomal rearrangements, and fusions in red indicate intrachromosomal rearrangement.

### Supplementary Tables

*Supplementary Table 1. Summary of nucleic acid extractions*

This table summarizes the quality assessment and quantitation of DNA and RNA extracted from the patient sample. These values include concentration and amount of RNA, RIN, and DV200 (quantified by the RNA Nano Chip on the Agilent Bioanalyzer 2100); the 260:280 ratio (quantified by Nanodrop); and the concentration and amount of DNA and DIN (quantified by the Genomic DNA Tape on the Agilent TapeStation 2200).

*Supplementary Table 2. Summary of sequencing libraries*

This table summarizes the quality assessment and quantitation of sequencing libraries, pre- and post-hybridized capture.
